# Multiple stressors in multiple species: effects of different RDX soil concentrations and differential water-resourcing on RDX fate, plant health, and plant survival

**DOI:** 10.1101/2020.05.21.108142

**Authors:** Richard F. Lance, Afrachanna D. Butler, Carina M. Jung, Denise L. Lindsay

## Abstract

Response to simultaneous stressors is an important facet of plant ecology and land management. In a greenhouse trial, we studied how eight plant species responded to single and combined effects of three RDX soil concentrations and two levels of water-resourcing. In an outdoor trial, we studied the effects of high RDX soil concentration and two levels of water-resourcing in three plant species. Multiple endpoints related to RDX fate, plant health, and plant survival were evaluated in both trials. Starting RDX concentration was the most frequent factor influencing all endpoints. Water-resourcing also had significant impacts, but in fewer cases. For most endpoints, significant interaction effects between RDX concentration and water-resourcing were observed for some species and treatments. Main and interaction effects were typically variable (significant in one treatment, but not in another; associated with increasing endpoint values for one treatment and/or with decreasing endpoint values in another). This complexity has implications for understanding how RDX and water-availability combine to impact plants, as well as for applications like phytoremediation. Two plant species native to the southeastern United States, *Ruellia caroliniensis and Salvia coccinea*, exhibited treatment responses that suggest they may be useful for phytoremediation, even within complex and changing environments.

## Introduction

One of the unique challenges to plant health on military ranges is environmental contamination with unique compounds required for military training and operations. One such compound is the nitroaromatic Royal Demolition Explosive (RDX; hexahydro-1,3,5-trinitro-1,3,5-triazine). Unignited RDX can leech into soil and groundwater and is a relatively common contaminant on military firing ranges [1-2]. Historically, RDX contamination of soils and waters also resulted from inadequate waste disposal practices at chemical and munitions production facilities [3-4]. RDX and its associated metabolites are absorbed by plant roots, then transported to stems, leaves, and flowers [5]. The majority of RDX may remain untransformed and accumulate in leaf tissues [3,6-7].

RDX soil contamination can be associated with negative trends in plant growth, survival, and health indicators, such as declines in chlorophyll concentrations or increased leaf chlorosis, necrosis, and/or curling [2, 7-10]. However, RDX can affect different species differently [10-12]. In a study of RDX impacts on 18 terrestrial plants [13], 16 species exhibited reduced growth and two exhibited enhanced growth at various RDX soil concentrations. Via et al. [14] noted that basic knowledge regarding the effects of RDX on wild plants and plant communities is lacking, including how RDX impacts different plant life stages. According to the same authors, additional research is needed to better understand how naturally occurring plant species respond to explosive contamination, and to better clarify the mechanisms involved in such responses.

In a period where natural resource managers, conservation professionals, and environmental planners are increasingly concerned about broad shifts in climate and changes in the frequencies of interannual climate extremes [15-18], interactions between climate and additional, other stressors are of particular interest. For example, air pollution can impair tree responses to freezing stress and drought conditions [19]. Water availability (e.g., annual precipitation) is one of several climatic factors for which both long-term shifts and short-term lability are of particular concern [20-22]. Water availability, whether in excess or deficiency, can stress plants. It also has the potential to influence the effects of RDX on plants, as transpiration significantly influences the movement of minerals, nutrients, and contaminants throughout a plant [23]. While interactions between climatic factors and RDX soil contamination are not well known, one study [8] found no interaction effect between water deprivation and RDX soil contamination on the production of anthocyanin and changes in leaf coloration in *Sida spinosa* L. (Malvaceae). Additional studies of a similar focus appear to be lacking, with more research needed. We report on the effects of different combinations of RDX soil concentration (50 ppm and 100 ppm) and water-resourcing levels to plant survival and health (leaf wilting, chlorosis, and chlorophyll content) for a total of nine different plant species. Limited RDX bioaccumulation data for each species under different treatments are also provided. The soil RDX concentrations used for these trials are well within those reported for RDX concentrations on contaminated sites [24].

## Materials and methods

### Greenhouse trial

#### Plant establishment and experimental treatments

For each of eight plant species (Table 1), 24-36 individual plants were potted in “clean” or RDX-infused soils and maintained in a greenhouse located on the US Army Engineer Research and Development Center’s (ERDC) Waterways Experiment Station (WES) in Vicksburg, MS. Soils were prepared using dry RDX (1% HMX; BAE System, Ordnance Systems Inc., Holston Army Ammunition Plant, Kingsport, TN) dissolved in acetone and mixed to target concentrations of 50 ppm and 100 ppm in a 10:3 silica quartz sand:loess soil (loess local to study area).

**Table 1.** Plant species and sample sizes per treatment (*n*_t_) for greenhouse studies.

| Species | Common Name | $n_t$ |
| --- | --- | --- |
| *Antirrhinum majus* | Snapdragon | 5 |
| *Dianthus* | Pink | 5 |
| *Hibiscus mocheutos* | Rose mallow | 5 |
| *Pentas lanceolata* | Starcluster | 5 |
| *Plumbago auriculata* | Cape leadwort | 2-3 |
| *Ruellia caroliniensis* | Wild petunia | 4 |
| *Salvia coccinea* | Scarlet Sage | 5 |
| *Tulbaghia violacea* | Society garlic | 2-4 |

Plants (n = 3–5) in each of the 0, 50, and 100 ppm soil concentration groups were further assigned to either a 1X or 0.5X water-resourcing treatment, for a total of six treatment groups (Table 2). Each plant received either approximately 1 L (1X) or 0.5 L (0.5X) of municipal tap water every other day. We assumed that at least one of the two levels of water-resourcing (1X or 0.5X) would constitute a stressor for each species, being comparatively further from the optimal water-resourcing level for that species under greenhouse conditions. Miracle-Gro^®^ Water Soluble All Purpose Plant Food (Scotts Miracle-Gro, Marysville, OH, USA) was included at an approximately 1:250 dilution with watering once per week. Plants were maintained in each treatment class for approximately 19 weeks (133 days).

**Table 2.** Water-resource and RDX soil concentration treatment classes employed in greenhouse trial.

| Water Resourcing Level \| RDX Soil Concentration Level |  |  |
| --- | --- | --- |
| 0.5X \| 0 ppm | 0.5X \| 50 ppm | 0.5X \| 100 ppm |
| 1X \| 0 ppm | 1X \| 50 ppm | 1X \| 100 ppm |

#### Measuring RDX concentrations in soil and plant tissues

RDX soil concentrations were measured at the start and end of trials, and in root tissues at trial completion. In all cases, RDX was extracted and measured using methods specified in EPA SW-846 Method 8330 [25]. Because of limited availability of above ground tissues, leaf and floral tissues, RDX concentration data for these tissues were not included in later analyses (though limited data for leaf and flower tissues are provided in Supporting Information, S1 File and S1 Table). Proportional reductions in soil RDX concentration (PRCs) for each unit (plant x species x treatment) were calculated by dividing the final soil RDX concentration in each plant (Supporting Information, S2 Table) by the overall mean soil RDX concentration for the treatment group (species x treatment), and then subtracting that value from 1. Bioconcentration factors (BCFs) for RDX in root tissues were calculated by dividing RDX concentrations in root tissue (Supporting Information, S2 Table) by the initial RDX concentrations in the soil.

#### Determining plant health endpoints

Upon approximately four weeks of growth under treatment conditions, three plant health endpoints (or stress metrics) were measured, namely the level of wilting observed on leaves, the level of chlorosis observed on leaves, and a chlorophyll content index (complete data found in Supporting Information, S3 Table). Wilting, or areas of dead tissue, was visually estimated for four randomly-selected leaves per plant. Each leaf was placed into one of four classes, based on estimated extent of wilted leaf surface area: 0–25%, 26–50%, 51–75%, 76–100%. A mean wilt level for each plant was then calculated. Mean leaf chlorosis, or areas of notably reduced or absent green hue, was estimated and calculated in the same fashion for each of the selected leaves. Additionally, for each selected leaf, one to three chlorophyll content index (*CCI*) measurements were made (depending on leaf surface area) using a CCM-200 plus chlorophyll content meter (Opti-Sciences, Inc.; Hudson, NH, USA). Either a single *CCI* value or mean leaf *CCI* was recorded for each leaf and mean 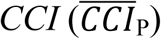 was calculated for each plant. The CCM-200 Plus uses two LED sources to measure light transmittance at two wavelengths: 931 nm, which falls within the chlorophyll absorbance range, and 653 nm, which provides a weighting factor associated with mechanical differences such as tissue thickness. *CCI* is the product of percent light transmittance at 931 nm and the inverse of the percent light transmittance at 653 nm. While not equivalent to actual density of chlorophyll in plant tissues, *CCI* provides a useful metric for comparing chlorophyll content among different samples.

### Outdoor plot trial

#### Outdoor plot establishment and experimental treatments

We selected three plant species, *P. lanceolata, R. caroliniensis*, and *Conradina canescens* for outdoor trials with single and combined water-level resourcing and RDX soil contamination treatments (Table 3). The former two species were included in this trial because they had provided a relatively consistent and productive bloom set in the greenhouse, and the outdoor trials were also part of a concurrent pollination study. The latter species, *C. canescens*, had thrived on the outdoor plot in an earlier pilot study. This species was thus included in this trial, despite being excluded from the earlier greenhouse trials due to rapid mortality across all treatments.

**Table 3.** Water-resource and RDX soil concentration treatment classes employed in outdoor plot trial.

| Water Resourcing Level \| RDX Soil Concentration Level |  |
| --- | --- |
| 0.5X \| 0 ppm | 0.5X \| 100 ppm |
| 1X \| 0 ppm | 1X \| 100 ppm |

Individual adult plants (n = 48 per species) were repotted into sandy Loess soil containing either no RDX (control) or 100 ppm RDX (prepared using methods described above). A section of a grassy field at ERDC-WES was developed as an outdoor study plot. The plot area was fenced to exclude large herbivores. Within the study plot, one individual potted plant from each species was placed into 35 L black rubber basin which had been placed on a plot “station” (Fig. 1). Each station was centered underneath a 0.9 m high clear polyethylene rain shield. A total of 48 stations were positioned in a grid with columns alternated between plants in control and RDX-contaminated soil. The first four rows of the grid was assigned to a 1X water-level treatment (approximately 1 L) while the second set of four rows was assigned to the 0.5X treatment (approximately 0.5 L). Plants were watered twice weekly. The plots was inspected for dead plants once per week for 10 weeks. After the end of this period, soil and shoot/leaf tissue were collected for measurement of RDX concentrations and calculation of PRCs and BCFs (as described previously; complete soil and shoot RDX concentration data available in Supporting Information, S4 Table).

**Figure 1.**
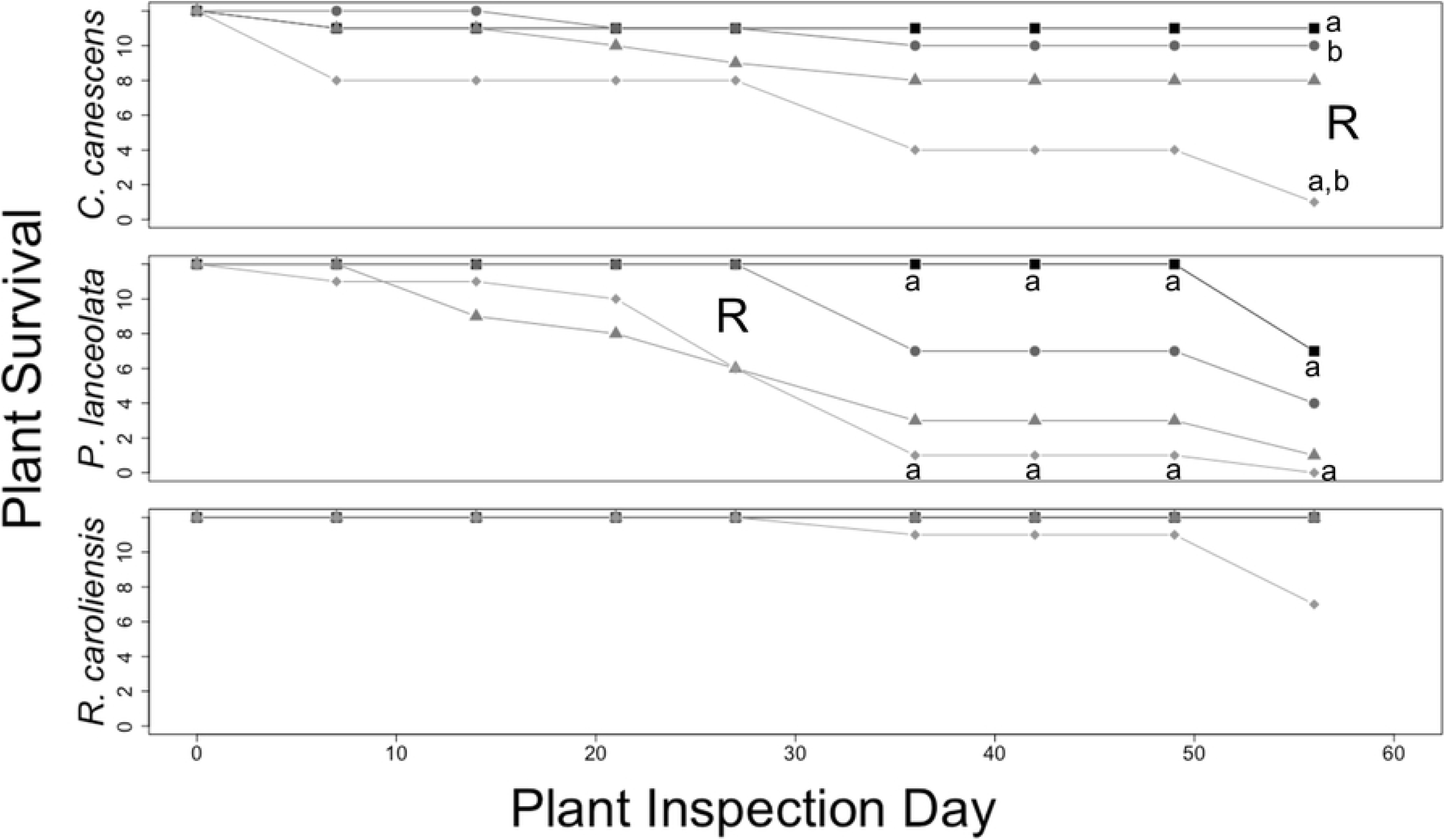
Images of outdoor plot at commencement of trial. *Above*, outdoor plot station with one potted plant per species for *R. caroliniensis* (left), *C. canescens* (upper right), and *P. lanceolata* (lower right). *Right*, outdoor experimental grid of 48 stations.

### Statistical analysis

For all analyses, because of relatively small sample sizes (n = 2–5), we set the critical value for assigning statistical significance at α = 0.10. The majority of treatment response datasets for all endpoints did not meet assumptions for parametric tests, so appropriate nonparametric tests were used. The effects of the main factors (water-resourcing levels, RDX soil concentration) and an interaction factor on each endpoint in the greenhouse trial (PRC, BCF, wilting, chlorosis, *CCI*) were tested using robust analysis of variance (RAOV [26-27]), provided in the package Rfit (version 0.24.2; [28]) in R (version 3.6.1 [29]). For each endpoint, pairwise comparisons of different treatment groups within each species were conducted using Dunn’s test [30], with a Bonferroni multiple comparisons correction to the critical value. Dunn’s tests were conducted using the R package FSA (version 0.8.25 [31]). The effect directions for significant factors in the various tests were assessed using interaction plots based on median values for each endpoint and rank-based estimates of regression coefficients [28].

For the outdoor plot trials, differences in PRC and BCF between plants of each species in the 0.5X|100 ppm and 1X|100 ppm treatment groups were assessed using one-sample Wilcoxon tests, performed in base R. For each species in the outdoor plot trial, differences in survival among treatments with different levels of the main factors or among plants in the different combined treatments (Table 3) within each species were assessed at each time point (plot inspection dates) using log-linear regression analysis, as calculated using the loglm function in the R package MASS (version 7.3-51.4 [32]). Pairwise comparisons of plant survival under different levels of each of the two main factors, or among the different combined treatments were assessed using chi-square tests with Bonferroni adjustments to critical values for multiple comparisons.

## Results and discussion

This study explored patterns in plant responses to an important military soil contaminant, the phytotoxic compound RDX, in the presence of an additional factor (and potential stressor), water-resourcing levels. We conducted tests with a total of nine plant species, none of which had previously been tested for responses to RDX soil contamination.

### Greenhouse trials: RDX uptake and plant health endpoint indices

We were able to maintain all eight plants species (Table 1) in all treatments in the greenhouse for the entire study period. However, because the 0.5X|100 ppm treatment group samples for *T. violacea* were spilled during sample processing and after remaining pot soils had been discarded, PRC could not be calculated for this treatment group in this species. Final soil PRCs (proportional reductions in RDX soil concentration) and root BCFs (bioconcentration factors) varied widely among the eight plant species and treatment groups (Figs. 2-3). In a few cases, soil samples had negative PRCs (and thus measured RDX concentrations above starting concentrations; Fig. 2). The initial soil RDX concentrations measured at the point of establishing plants in pots (time point 0) had *mean* RDX concentrations of 50 ppm and 100 ppm (data not shown), and individual measures exceeding these mean values at the end of the trial likely represent, to some degree, nonuniform mixing of RDX in soil and associated random sampling error.

**Figure 2.**
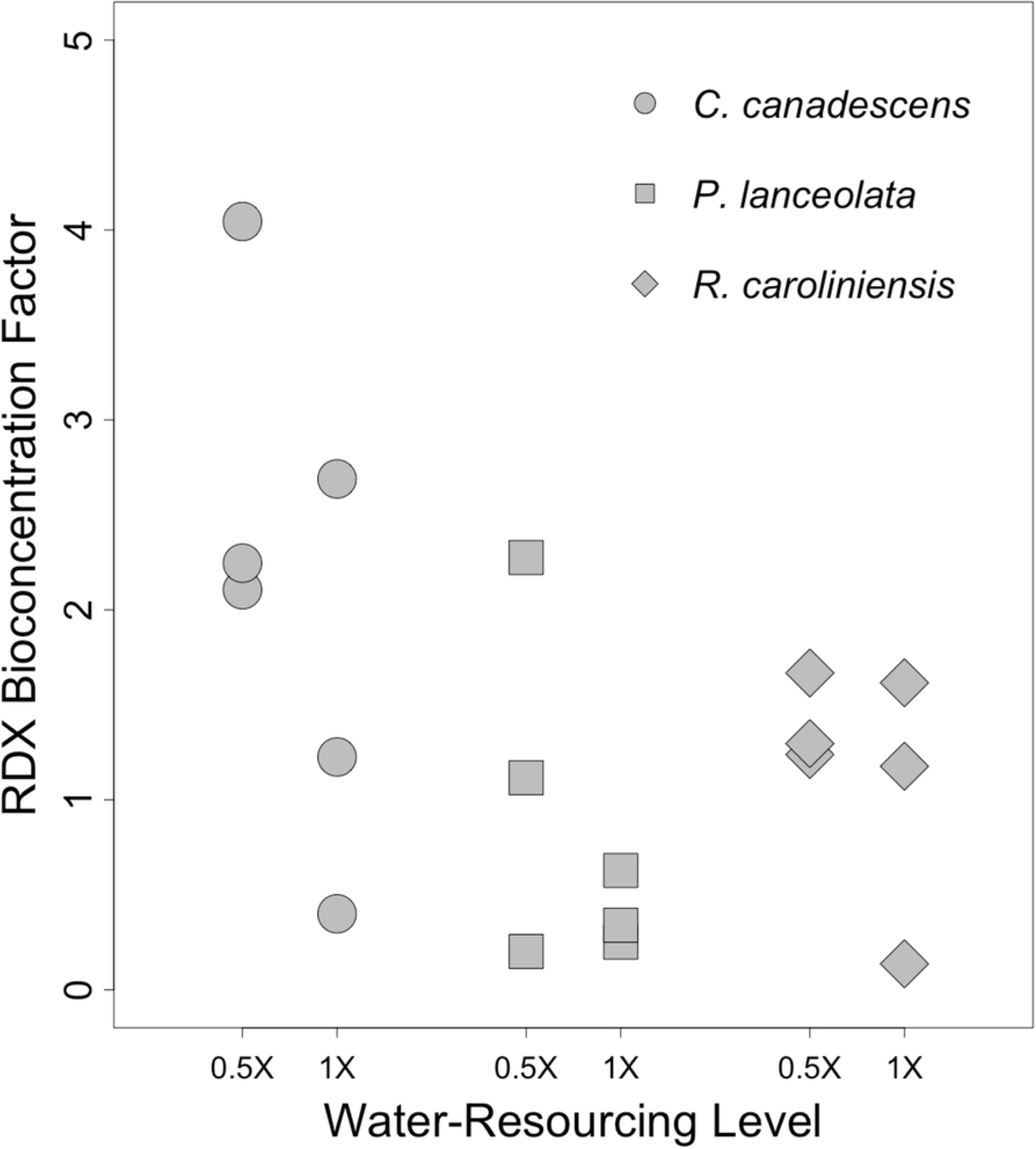
Proportional reductions in soil RDX in greenhouse trials. Boxplots of proportional reductions in *soil* RDX concentrations (PRCs) for eight plant species reared for 133 days in a greenhouse under different combinations of two factors, water-resourcing level (1X and 0.5X) and soil RDX concentration (50 and 100 ppm). Boxplots include median (bold horizontal line) and mean (dotted horizontal line) values. Sample sizes within each group are found on the x-axis. R = significant effect of soil RDX concentration; W = significant effect of water-resourcing level; I = significant interaction effect; a,b = significant pairwise difference between treatment groups. α ≤ 0.10.

**Figure 3.**
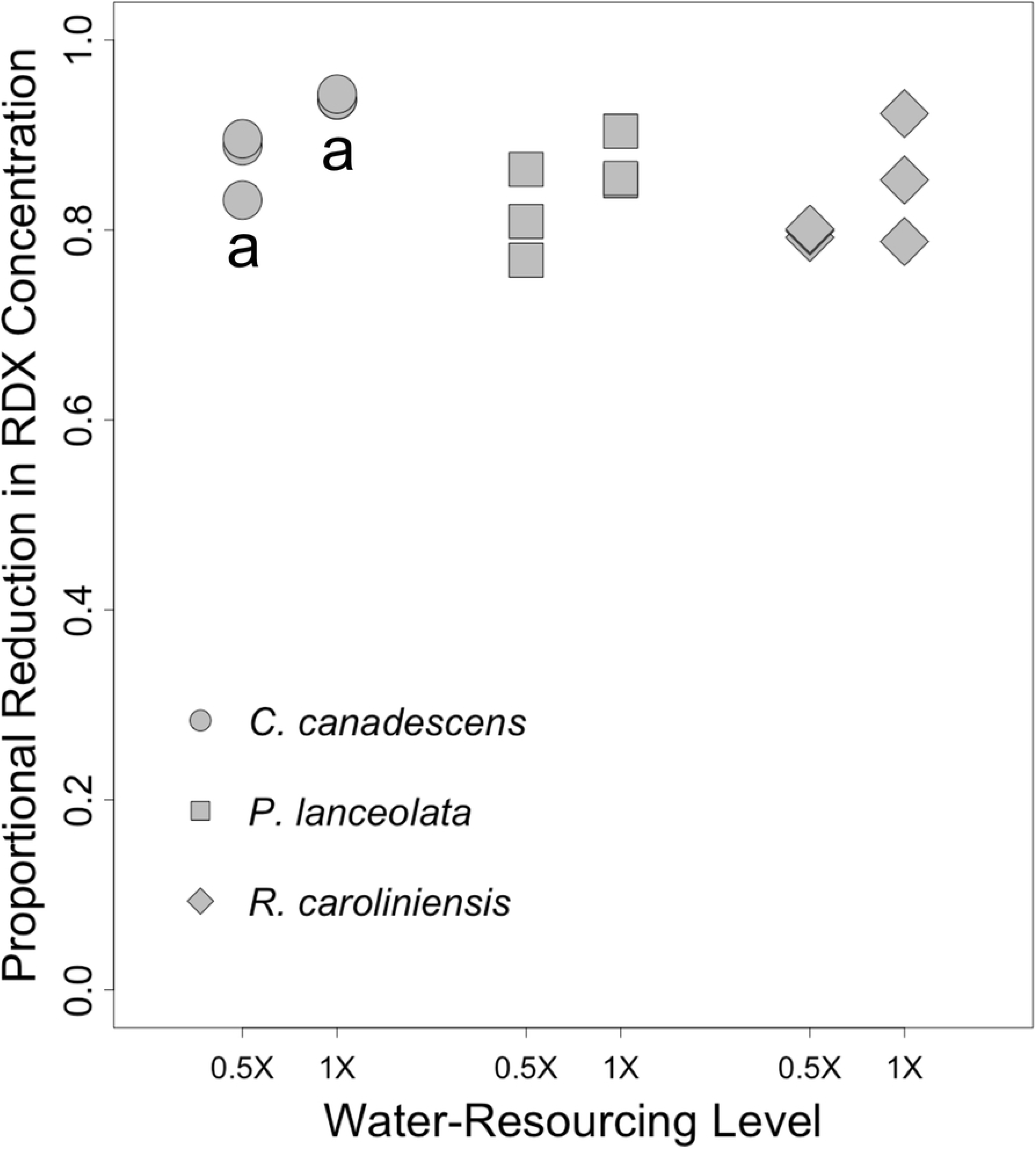
RDX bioconcentration factors in greenhouse trials. Boxplots of final *root* RDX bioconcentrations factors (BCFs) for eight plant species reared for 133 days in a greenhouse under different combinations of two factors, water-resourcing level (1X and 0.5X) and soil RDX concentration (50 and 100 ppm). Boxplots include median (bold horizontal line) and mean (dotted horizontal line) values. Sample sizes within each group are found along the top of each boxplot frame. R = significant effect of soil RDX concentration; W = significant effect of water-resourcing level; I = significant interaction effect; I = significant interaction effect; a = significant pairwise difference between treatment groups. α ≤ 0.10.

RDX soil concentrations were a significant factor in PRC and root BCF for a majority of species (Figs. 2-3). In many cases, PRC values for the 100 ppm RDX treatments were higher than with the 50 ppm RDX treatments, but pairwise comparisons were not always statistically significant (Figs. 2). In terms of bioaccumulation, the 50 ppm RDX treatments generally exhibited higher root BCF values than treatments with 100 ppm RDX, though, again, pairwise comparisons were not always statistically significant (Fig. 3). As a single factor, water-resourcing had little effect on PRC across species and treatments (Fig. 2), but was more commonly a factor in root BCF (Fig. 3). In three plant species, reduced water-resourcing was associated with lower root BCFs (Fig. 3). Of particular interest, significant interaction effects between the main factors (RDX soil concentrations and water-resourcing levels) were apparent for both PRCs and BCFs within several species. The 0.5X|100 ppm treatment resulted in the highest levels of PRC for most species, though PRCs for this treatment group was not always statistically different from other treatment groups (Fig. 2). In terms of root RDX accumulation, the 01X |100 ppm treatment group often exhibited the highest BCFs (Fig. 3). Significant interaction effects were observed for both PRCs and BCFS, in three and two species respectively (Figs. 2-3). The strength (significant or not significant) and direction (increasing or decreasing endpoint values) of those effects varied from case to case (Table 4).

**Table 4.**
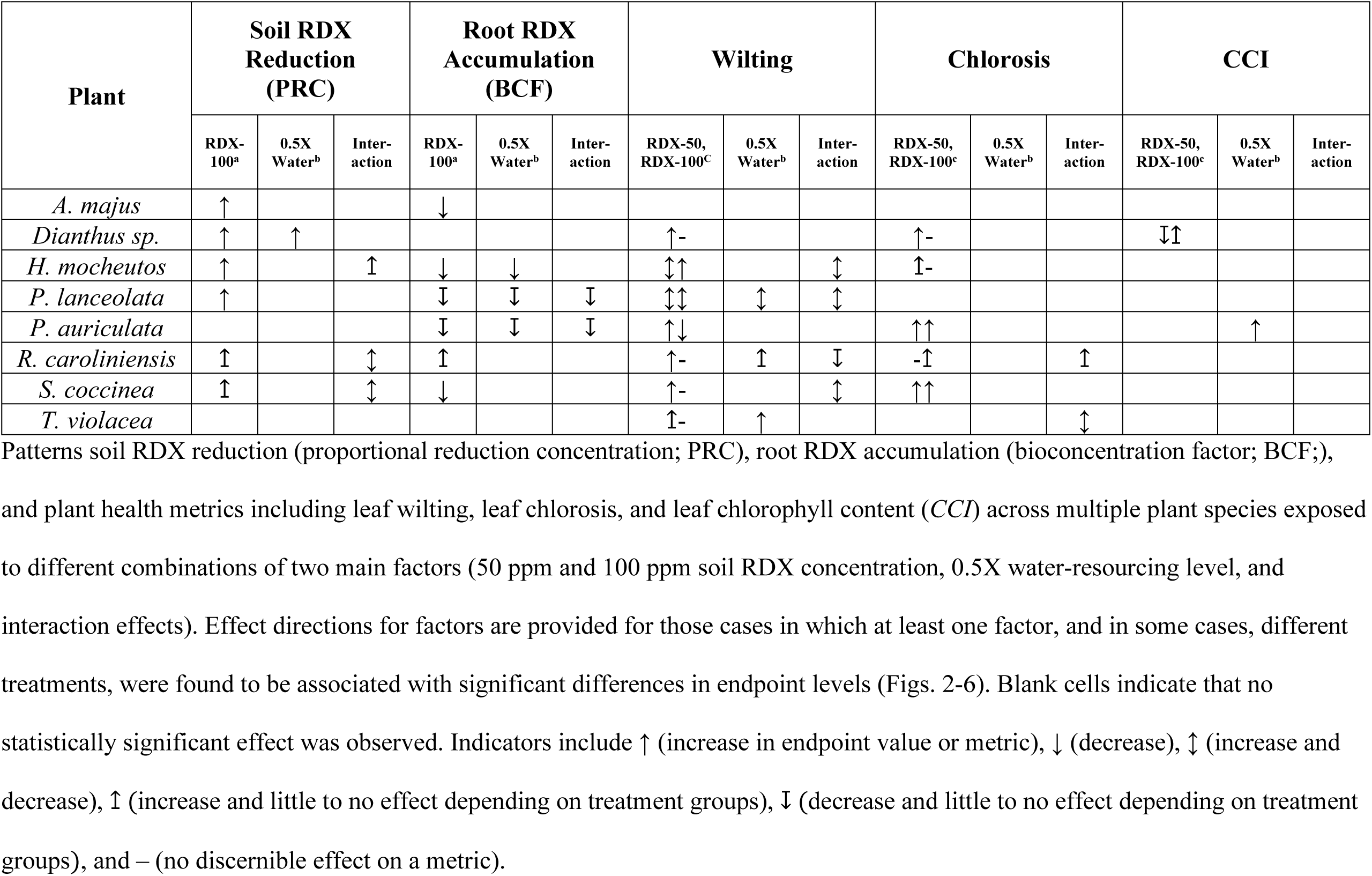
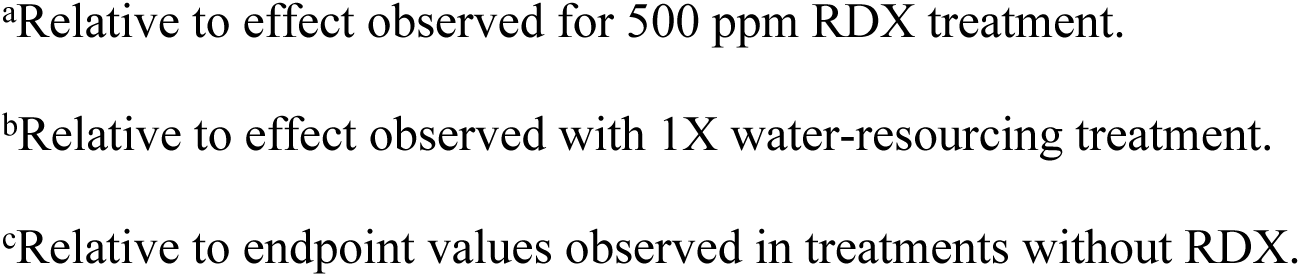
Summary of patterns in RDX fate and plant health in greenhouse trial.

Patterns soil RDX reduction (proportional reduction concentration; PRC), root RDX accumulation (bioconcentration factor; BCF;), and plant health metrics including leaf wilting, leaf chlorosis, and leaf chlorophyll content (*CCI*) across multiple plant species exposed to different combinations of two main factors (50 ppm and 100 ppm soil RDX concentration, 0.5X water-resourcing level, and interaction effects). Effect directions for factors are provided for those cases in which at least one factor, and in some cases, different treatments, were found to be associated with significant differences in endpoint levels (Figs. 2-6). Blank cells indicate that no statistically significant effect was observed. Indicators include ↑ (increase in endpoint value or metric), ↓ (decrease), ↕ (increase and decrease), ↥ (increase and little to no effect depending on treatment groups), ↧ (decrease and little to no effect depending on treatment groups), and – (no discernible effect on a metric).

**Figure 4.**
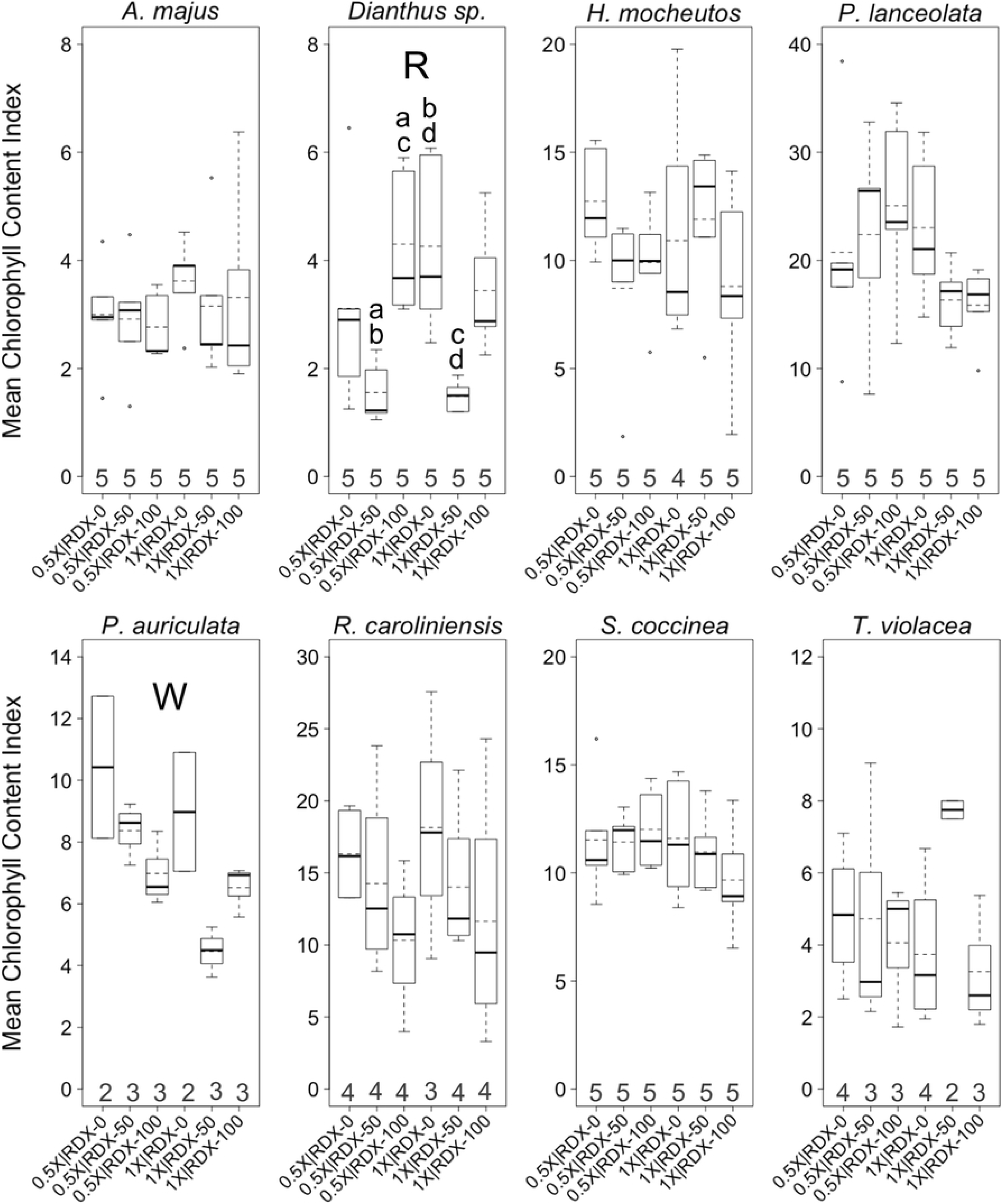
Wilting levels in greenhouse trials. Boxplots of mean *wilting* levels per plant across eight plant species. Plants were reared in a greenhouse under different combinations of two factors, water-resourcing level (1X and 0.5X) and soil RDX concentration (0, 50, and 100 ppm). Boxplots include median (bold horizontal line) and mean (dotted horizontal line) values. Sample sizes within each group are found along the top of each boxplot frame. R = significant effect of soil RDX concentration; W = significant effect of water-resourcing level; I = significant interaction effect; a–d = significant pairwise difference between treatment groups. α ≤ 0.10.

**Figure 5.**
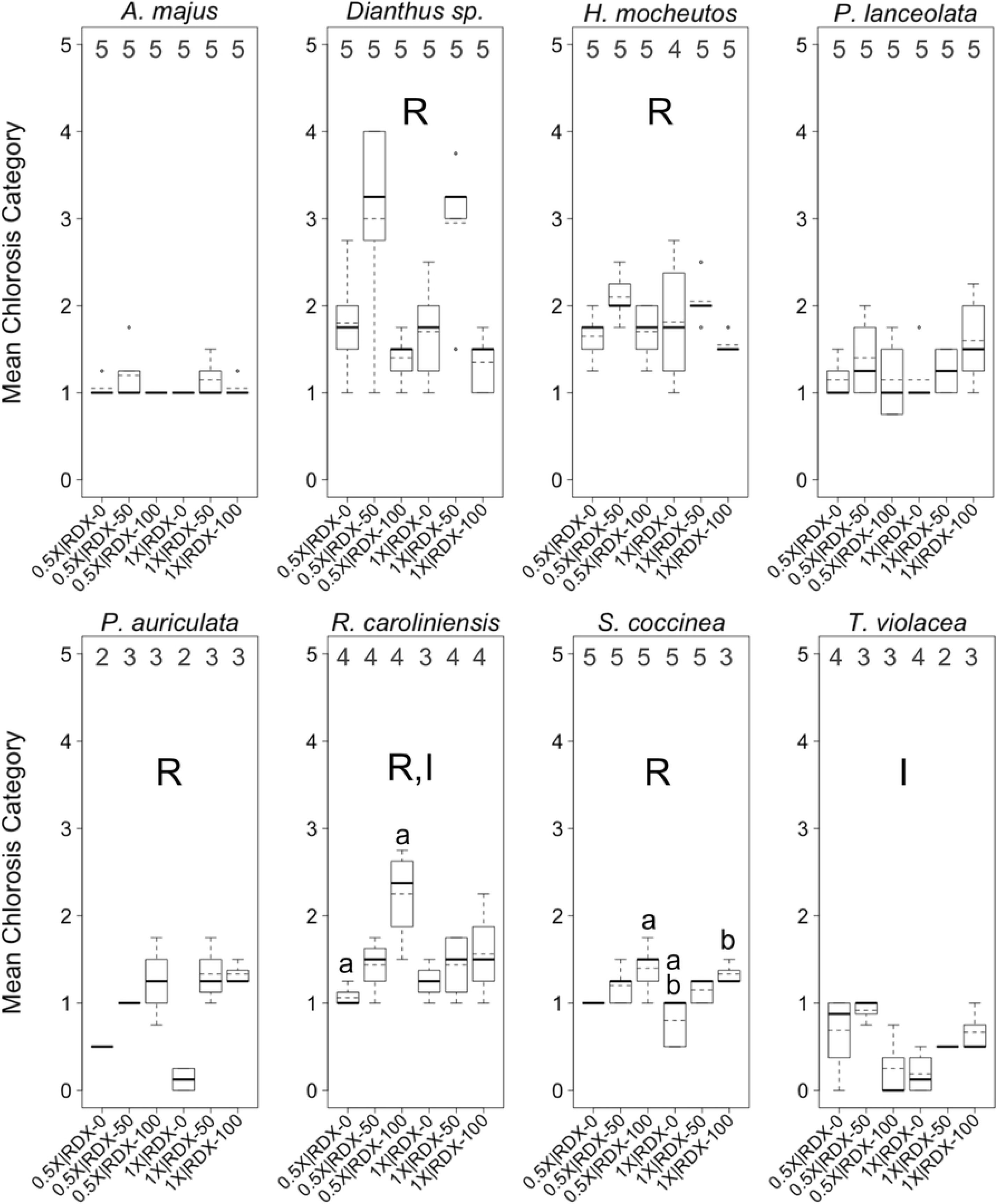
Chlorosis levels in greenhouse trials. Boxplots of mean *chlorosis* levels per plant across eight plant species. Plants were reared in a greenhouse under different combinations of two factors, water-resourcing level (1X and 0.5X) and soil RDX concentration (0, 50, and 100 ppm). Boxplots include median (bold horizontal line) and mean (dotted horizontal line) values. Sample sizes within each group are found along the top of each boxplot frame. R = significant effect of soil RDX concentration; I = significant interaction effect; a–d = significant pairwise difference between treatment groups. α ≤ 0.10.

**Figure 6.**
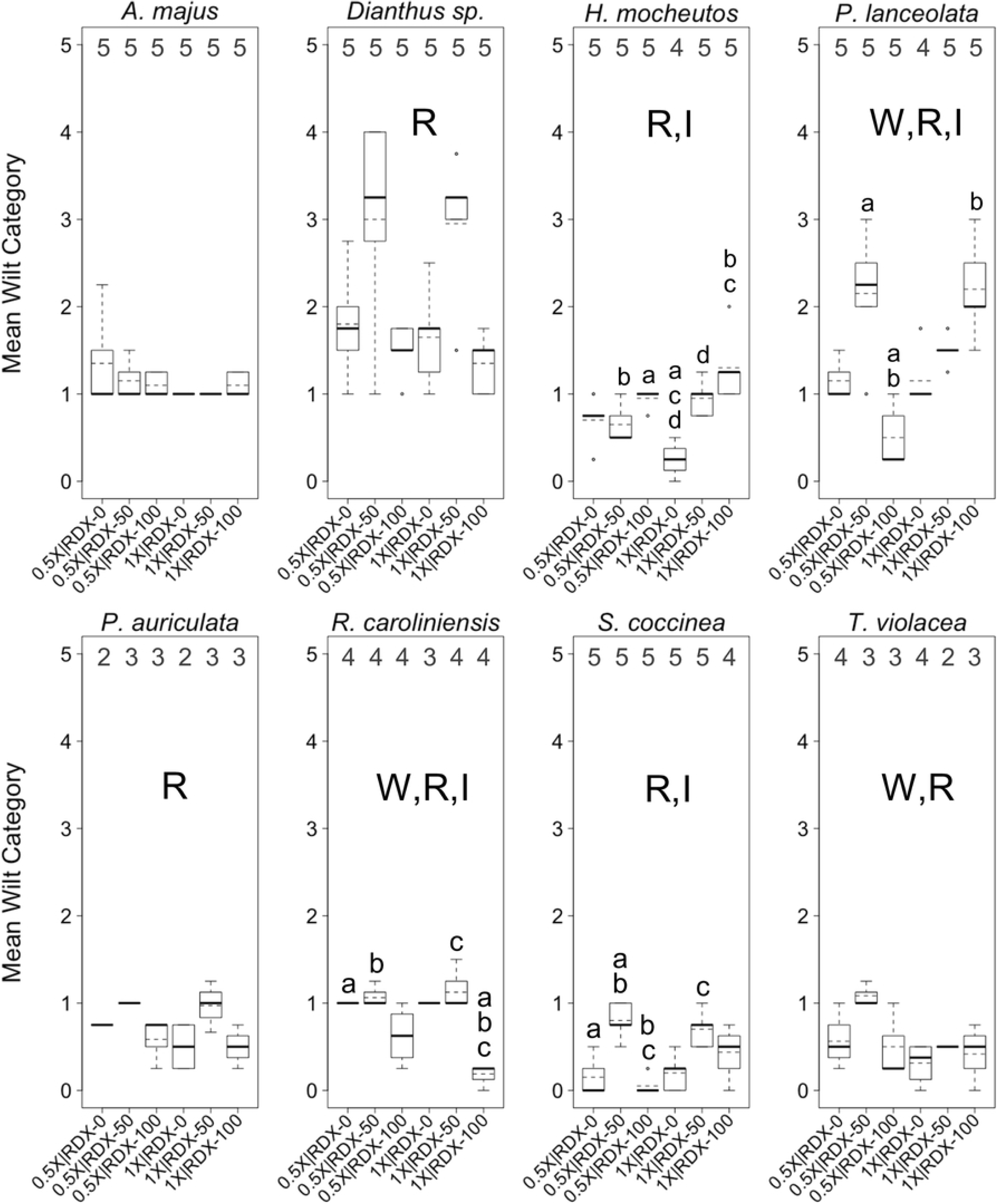
Chlorophyll content index levels in greenhouse trials. Boxplots of mean *chlorophyll content index* (*CCI*) levels per plant across eight plant species. Plants were reared in a greenhouse under different combinations of two factors, water-resourcing level (1X and 0.5X) and soil RDX concentration (0, 50, and 100 ppm). Boxplots include median (bold horizontal line) and mean (dotted horizontal line) values. Sample sizes within each group are found on the x-axis. R = significant effect of soil RDX concentration; W = significant effect of water-resourcing level; a–d = significant pairwise difference between treatment groups. α ≤ 0.10.

The effects of different water-resourcing levels, soil RDX concentrations, and the interaction of these factors on wilting levels were varied and complex for nearly all of the eight plant species within the greenhouse trials (Fig. 4). One general trend was that different RDX concentrations were most common factor influencing plant health responses to different treatments (Fig. 4-6; Table 4). Most plants across treatments and species exhibited only mild wilt (i.e., 0–25%), or in several cases, moderate wilt (i.e., 25–50%; Fig. 3). Higher degrees of wilt (> 50% or greater of leaf surface) were relatively rare, and were observed in only a few treatment groups within *Dianthus* and *P. lanceolata*. In seven of eight species, there was a statistically significant association between starting RDX soil concentrations and observed differences in mean wilting category. In some cases, higher RDX concentrations appeared to be associated with increasing levels of wilt, yet conversely, in some cases, higher soil RDX concentrations appeared to benefit plants (reduced levels of wilt; Fig. 4; Table 4). Differences in water-resourcing levels were associated with significant differences in mean wilting category in three species, with varied effect strengths and directions (Fig. 4; Table 4). Significant interaction effects between the main factors were observed in four species, with the strength and direction of those effects being mixed, even within species (Fig. 4; Table 4).

Most plants, across treatments and species, exhibited moderate levels of leaf chlorosis (i.e., 26–50%), with relatively few observed cases of milder (i.e., 0–25%) or more pronounced (51–100%) chlorosis (Fig. 5). *Dianthus* exhibited somewhat higher chlorosis levels than other species, while *T. violacea* exhibited notably lower levels of chlorosis. As with wilting, RDX soil concentration was, by far, the most common factor in significant differences in chlorosis levels among treatments. In general, RDX soil contamination increased levels of chlorosis, with little difference in the effects of the two different RDX concentrations. Water-resourcing did not appear to influence levels of chlorosis. Of particular interest, significant interaction effects were observed in *R. caroliniensis* and *T. violacea*. In both cases, the strength or direction of the interaction effect on chlorosis was mixed.

In addition to experimentally testing for effects of different dual-factor treatments on common plant health indices like wilting and chlorosis, a distinctive feature of our study included an attempt to gauge the impacts of these treatments on chlorophyll levels in leaf tissues. This was done through measurements of a chlorophyll context index (*CCI*). Though similar metrics have been used in a few other RDX phytotoxicity studies [2, 33], to the best of our knowledge, *CCI* has not been previously used to gauge the plant health impacts of RDX or other munitions.

*CCI* varied widely between plants, but there were very few significant differences among factors or treatment groups (Fig. 6). RDX soil concentration was a factor associated with significant differences in 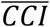 for one species (*Dianthus* sp.), as was water-resourcing (*P. auriculata*). Interestingly, the lower water-resourcing level was associated with higher 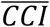 in *P*.

*auriculata* (Fig. 6). There were no other significant associations between this water-resourcing level and relatively better health in this species, though a larger sample size may have supported an association with a reduced level of chlorosis (Figs. 4,5). It would be tempting to assume that the 0.5X water-resourcing level was optimal for *P. auriculate*, however the 1X|0 ppm treatment groups exhibited the lowest (or among the lowest) levels of wilting and chlorosis within this species. There were no significant factor interaction effects on 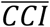 for any of the tested species.

#### Perspectives from greenhouse trial

The dynamics of RDX uptake, root bioaccumulation, and plant health within RDX contaminated sites are clearly complex. Predictions regarding soil contaminant uptake and fate in plants, as well as predicted plant health responses to RDX soil contamination, will, not unexpectedly, be insufficient when based on a single factor. The trend for greater proportional reduction in soil RDX when RDX concentrations are higher (Fig. 2), with some interaction effect associated water availability, has implications for phytoremediation practices, and, perhaps, particularly for phytoremediation in regions that may experience reduced or more variable precipitation in the future. The fact that PRCs in the southeastern U.S. native plant *P. auriculata* were notably consistent, regardless of starting RDX concentrations in the soil or water-resourcing, make this plant a promising candidate for phytoremediation efforts in that region.

The high inter-variability in health effects observed across species and between treatments also reinforces that benefits to phytoremediation efforts of local pilot studies. Such studies would help overcome some of the inherent unpredictability of how plants will respond to different combinations of environmental factors. The fact that the southeastern U.S. native plant *S. coccinea* exhibited some of the lowest wilting and chlorosis levels, and consistent 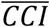 levels, regardless of starting RDX concentrations in the soil or water-resourcing, make this species another promising candidate for phytoremediation efforts in that region.

### Outdoor plot trial: RDX concentrations and plant survival

As evidenced by PRC levels, at the end of the outdoor trial, RDX soil concentrations were much lower than the starting concentration of 100 ppm (Fig. 7). PRCs were not widely divergent among species (Fig. 7). Though there appeared to be a trend towards higher PRCs associated with 1X water-resourcing, this pattern was only statistically significant in the case of *C. canescens* (*p* = 0.10).

**Figure 7.**
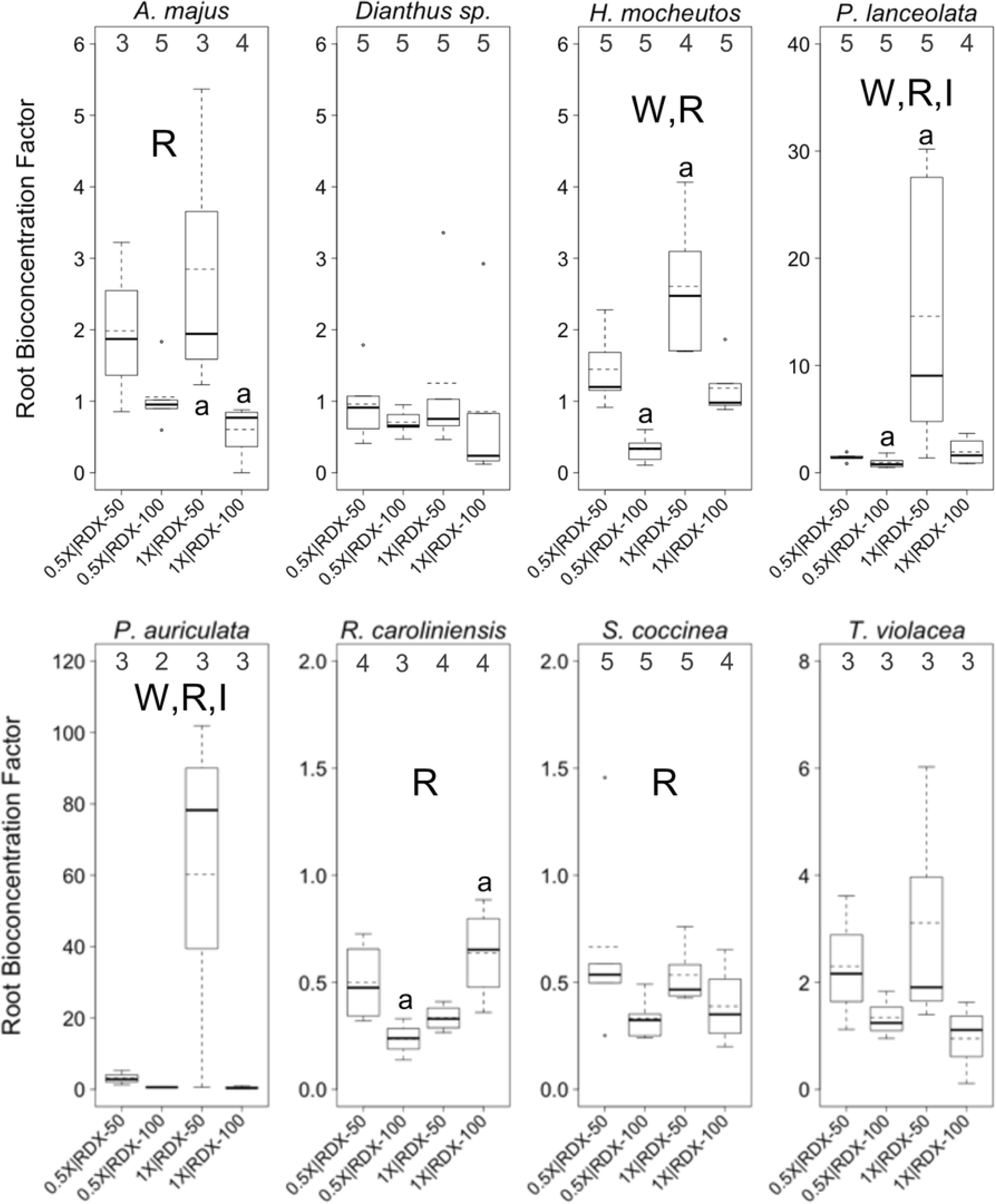
Proportional reductions in soil RDX in outdoor trials. Proportional reductions concentrations (PRCs) in soils from pots containing three different plant species maintained outdoors with either a 1X or 0.5X water-resourcing treatment for at least 41 days. Initial concentration of RDX in potting soils was 100 ppm. For each species, soils from n = 3 plants were sampled per treatment group. a = significant pairwise difference between treatment groups (*p* ≤ 0.10).

By the end of the trial, plants from all three species had bioaccumulated RDX. In most cases, BCFs were greater than the starting concentration of 100 ppm (BCF > 1). Though there appeared to be a trend of higher BCFs within the 0.5X water-resourcing treatment, there were no statistically significant differences among treatment groups.

All three plant species exhibited different survival patterns, with RDX being the primary factor associated with plant mortality (Fig. 9). In *C. canescens*, plant survival significantly diverged between treatments with and without RDX by Day 56 (*p* = 0.029), and *P. lanceolata* by Day 26 (and persisting through Day 56 (*p* = 0.002-0.046)). There was, to an extent, a combined effect between RDX concentrations and water-resourcing, as the 0.5X|100 ppm treatment group consistently exhibited the lowest survival rate. This treatment group exhibited significantly different survival from several other treatment groups at Day 56 in *C. canescens* (*p* = 0.004-0.020), and from the 1X|0 ppm treatment group in *P. lanceolata* starting on Day 26 (and persisting through Day 56 (*p* = 0.002-0.046)). For *R. caroliniensis*, only the 0.5X|100 ppm treatment group exhibited any plant mortality, beginning between Day 27 and Day 36. There were no statistically significant differences in plant survival among treatment groups in this species.

**Figure 8.**
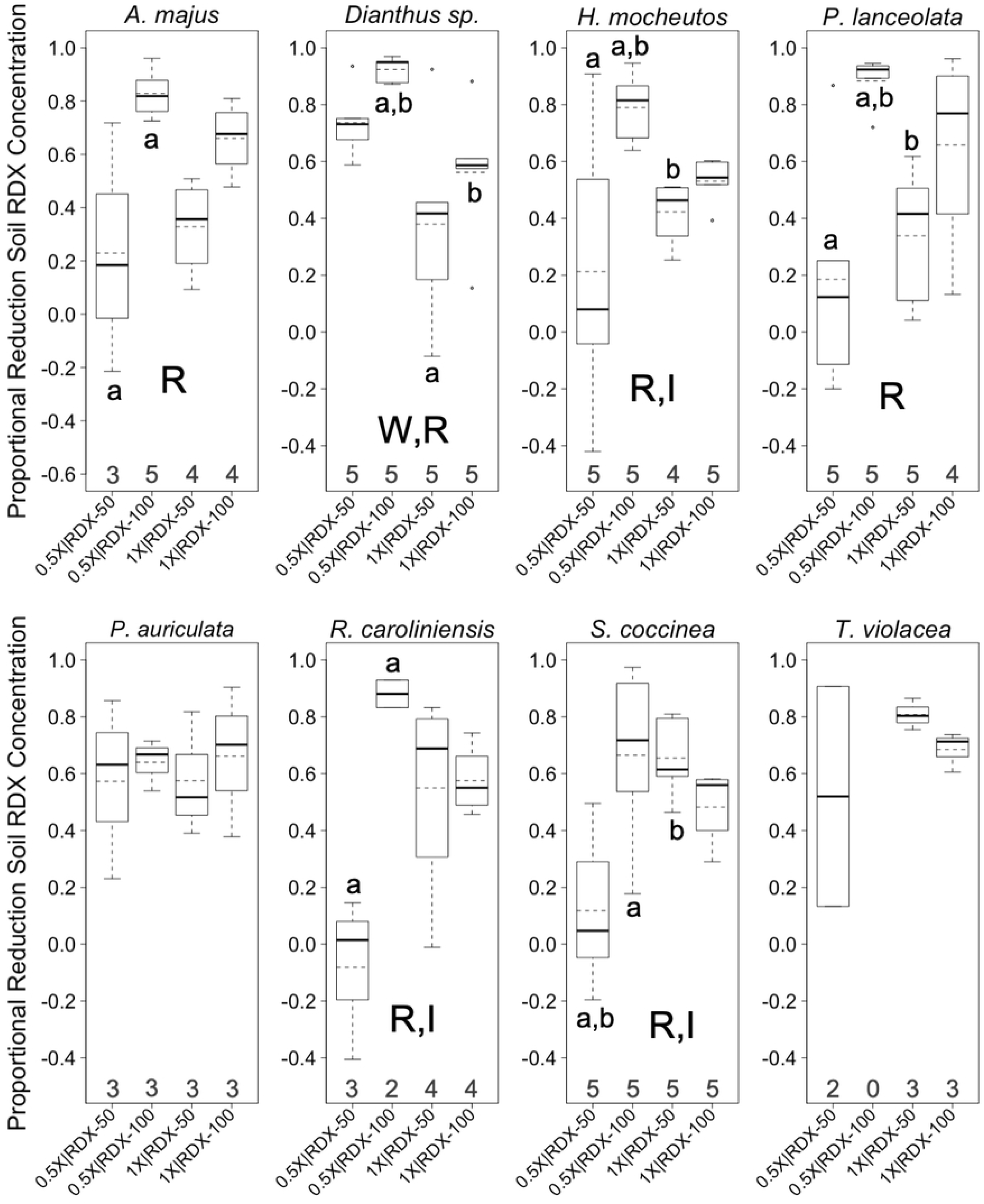
RDX bioconcentration factors in outdoor trials. Bioconcentration factors (BCFs) for RDX in leaf and shoot tissues of three different plant species maintained outdoors with either a 1X or 0.5X water-resourcing treatment for at least 41 days. Initial concentration of RDX in potting soils was 100 ppm. For each species, above-ground tissues from n = 3 plants were sampled per treatment group.

**Figure 9.**
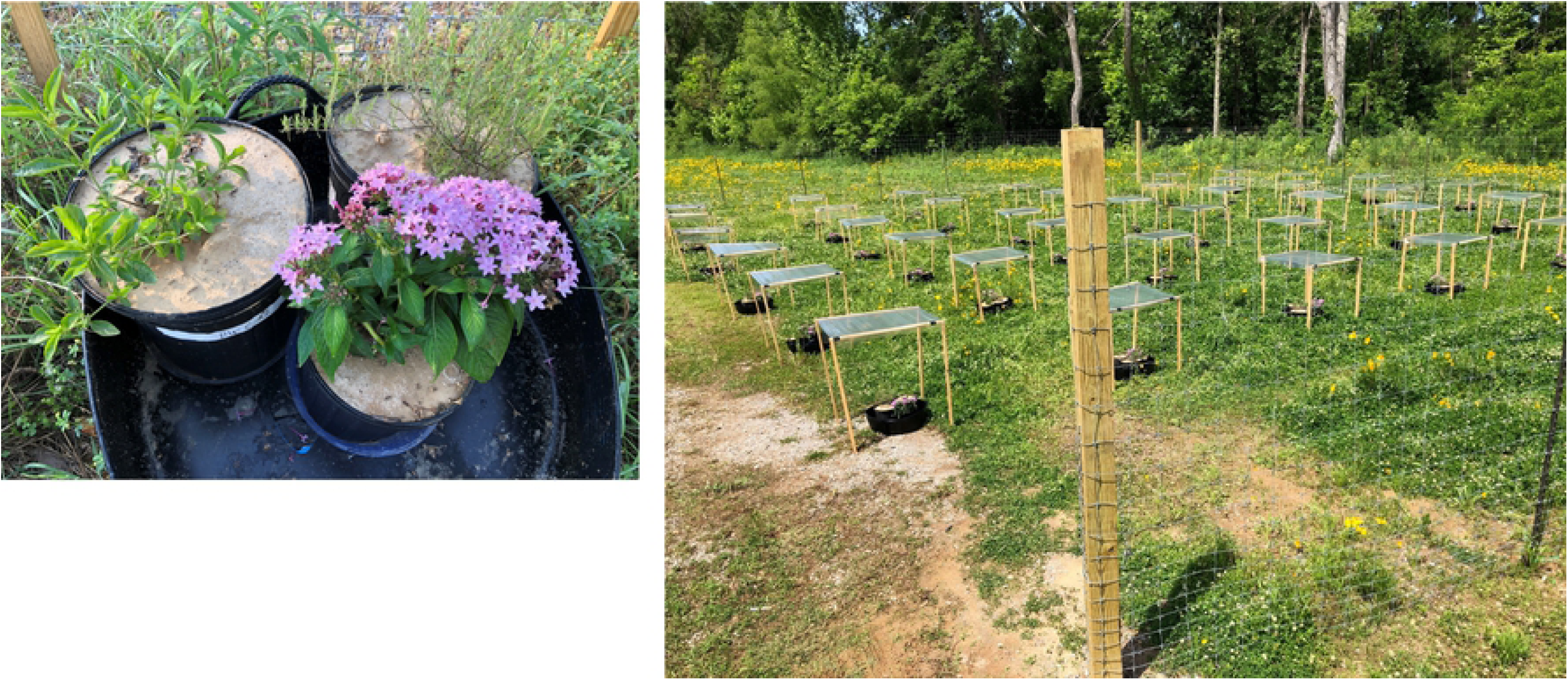
Plant survival in outdoor plot trial. Numbers of surviving *C. canescens, P. lanceolata, and R. caroliniensis* under four treatments over 56 days. Treatments included reduced water-resourcing (1X vs. 0.5X) and soil contamination with RDX (0 vs 100 ppm). R = day by which significant differences in survival had emerged (adjusted α = 0.05); a–b = day by which significant pairwise difference between treatment groups (designated by same letter) emerged (adjusted α = 0.017).

#### Perspectives from outdoor plot trial

RDX uptake and bioaccumulation of plants under approximately natural conditions appeared to be influenced by water availability, though without stronger statistical patterns and additional, more comprehensive data (e.g., root BCF, changes in plant biomass), a greater understanding of the mechanisms involved is not possible. As was the case with plant health in the greenhouse trials, initial RDX concentrations was the more influential factor in plant survival on outdoor plots than water-resourcing. Also, as observed with plant health, soil contamination and water availability appeared to have an interaction effect on plant survival in some cases (*C. canescens*, and, perhaps, *R. caroliniensis*), but not all (*P. lanceolata*). This highlights the fact that plant survival in RDX contaminated soil is likely a complex and difficult-to-predict phenomenon, influenced by additional factors or stressors. The southeastern U.S. native plant *R. caroliniensis*, which, in greenhouse trials, exhibited promising attributes relative to use in phytoremediation, also exhibited substantially lower mortality than the other two test species, despite similar RDX uptake and bioconcentration levels.

## Conclusions

In this study we demonstrated RDX uptake and bioconcentration in nine previously untested species of plants. We further experimentally demonstrated that an additional factor, water-resourcing, could significantly change plant-RDX interactions. In some cases, interaction effects between the two main factors (soil RDX, water-resourcing) emerged. The impacts of soil RDX, water-resourcing, and interaction effects of these two factors on plant RDX uptake, bioconcentration, health, and survival were typically complex and not easily generalizable. These observations have implications for understanding how plant species (and hence, plant communities) might respond to RDX soil contamination under different climatic scenarios, and for selecting specific plant species for phytoremediation of RDX.

## Acknowledgements

We are grateful for support from the U.S. Army Environmental Quality and Installations Program leadership and staff. We express gratitude for N. Harms, X. Guan, J. Smith, C. Roesch, K. Holmes, E. Lance, J. Lance, M. Lance, B. Jung, M. Carr, B. DeRossette, T. O’Neil, and M. Ogburn for assistance in setting-up and maintaining greenhouse facilities and the outdoor plot. We also thank M. Beverly, G. Watson, R. Jarratt, T. O’Neal, T. Nabors, A. Kessee, S. Zetterholm, and T. Roberts for assistance in greenhouse data collection and processing. We are grateful to K. Indest, C. Thomas, and S. Feist for insightful manuscript reviews. Views, opinions and/or findings contained herein are those of the authors and should not be construed as an Official Department of the Army position or decision unless so designated by other official documentation.

## Supporting information

**S1 File. RDX concentrations for leaf and flower tissues in greenhouse trials**. A brief description of methods and results for RDX concentrations in several plant speciess maintained under different treatments in the greenhouse trial.

**S1 Tables A and B. Leaf and flower RDX concentration data from greenhouse trial**. Tables of RDX concentrations for soil (Table A, “soil_rdx”) and leaves (Table B, “leaf_rdx”) for several plant species (“plant_species”) maintained under different treatments (“treatment”) in the greenhouse trial. Treatment groups were based on different initial soil concentrations of RDX (“rdx”) and water-resourcing (“water”).

**S2 Table. Soil and root RDX concentration data from greenhouse trial**. Table of soil and root RDX concentrations (“soil_rdx,” “root_rdx,” respectively) for each individual plant (“unit”) within each treatment group (“treatment”) and within each of eight species (“plant_species”; Table 1). Treatment groups based on different initial soil concentrations of RDX (“rdx”) and water-resourcing (“water”).

**S3 Tables A, B, and C. Wilting, chlorosis, and chlorophyll content index data from greenhouse trial**. Tables of estimated wilting levels (Table A), estimated chlorosis levels (Table B), and chlorophyll content index values (Table C) for four leaves (M1-M4) from each inidividual plant (“unit”) within each treatment group (“treatment”) and from within each of eight species (“species”; Table 1). Treatment groups were based on different initial soil concentrations of RDX (“rdx”) and water-resourcing (“water”).

**S4 Tables A and B. Soil and shoot RDX concentration data from outdoor plot trial**. Tables of soil (Table A) and shoot (Table B) RDX concentrations (“soil_rdx,” “shoot_rdx,” respectively) for individual plants within each treatment group (“treatment”) and within each of three species (“plant_species”; Table 1). Treatment groups were based on different initial soil concentrations of RDX (“rdx”) and water-resourcing (“water”).

